# Discovering hidden brain network responses to naturalistic stimuli via tensor component analysis of multi-subject fMRI data

**DOI:** 10.1101/2021.01.14.426756

**Authors:** Guoqiang Hu, Huanjie Li, Wei Zhao, Yuxing Hao, Zonglei Bai, Lisa D. Nickerson, Fengyu Cong

## Abstract

The study of brain network interactions during naturalistic stimuli facilitates a deeper understanding of human brain function. Intersubject correlation (ISC) analysis of functional magnetic resonance imaging (fMRI) data is a widely used method that can measure neural responses to naturalistic stimuli that are consistent across subjects. However, interdependent correlation values in ISC artificially inflated the degrees of freedom, which hinders the investigation of individual differences. Besides, the existing ISC model mainly focus on similarities between subjects but fails to distinguish neural responses to different stimuli features. To estimate large-scale brain networks evoked with naturalistic stimuli, we propose a novel analytic framework to characterize shared spatio-temporal patterns across subjects in a purely data-driven manner. In the framework, a third-order tensor is constructed from the timeseries extracted from all brain regions from a given parcellation, for all participants, with modes of the tensor corresponding to spatial distribution, time series and participants. Tensor component analysis (TCA) will then reveal spatially and temporally shared components, i.e., naturalistic stimuli evoked networks, their temporal courses of activity and subject loadings of each component. To enhance the reproducibility of the estimation with TCA, a novel spectral clustering method, tensor spectral clustering, was proposed and applied to evaluate the stability of TCA algorithm. We demonstrate the effectiveness of the proposed framework via simulations and real fMRI data collected during a motor task with a traditional fMRI study design. We also apply the proposed framework to fMRI data collected during passive movie watching to illustrate how reproducible brain networks are identified evoked by naturalistic movie viewing.

## Introduction

There is growing interest in studying brain function during naturalistic stimuli e.g. film clips, spoken narratives and music as experimental paradigms to investigate human cognition and behavior in “real-world” (Breakspear and Chang, 2020; Hasson et al., 2004; Huth et al., 2016; Nishimoto et al., 2011; Sonkusare et al., 2019; Spiers and Maguire, 2007). Naturalistic stimulus viewing during functional magnetic resonance imaging (fMRI) is emerging as a powerful tool to define brain imaging-based markers of psychiatric illness (Eickhoff et al., 2020), with several advantages in comparison to unconstrained resting state. Namely, studying brain network function during naturalistic stimulus viewing may facilitate a deeper understanding of human brain function since the passive state is better constrained and it reduces participant motion, which greatly increases the quality of the fMRI data. However, new analytical strategies are needed that assess both the shared rapid temporally evolving brain responses evoked by the naturalistic stimuli in participants, as well as the idiosyncrasy in individual participants (Simony and Chang, 2020).

The most popular approach for analysis of naturalistic fMRI data is assessment of intersubject correlations (ISC), which quantifies the across-subject consistency of stimulus-driven responses (Hasson et al., 2004). While evoked brain activity using traditional fMRI study designs and stimuli is relatively straightforward to model, naturalistic stimuli are complex and dynamic, and it is much more difficult to generate a model of evoked activity for analyses. Instead, for dynamic complex stimuli such as movies, ISC measures shared information across brains by using each individual’s brain activity measured by fMRI to model another individual’s brain activity. Using this strategy, the shared brain regions that respond to the same time-locked naturalistic stimuli across subjects can be estimated, even with stimuli that reflect complex dynamic real-life contexts (Hasson et al., 2004; Kauppi et al., 2014; Nastase et al., 2019). Modifications to intersubject functional correlation (IFC), temporal intersubject functional correlation (ISFC) has been proposed that consider the timecourse correlations between all possible pair-wise combinations of brain parcels across subjects (Nastase et al., 2019) and spatial ISC is an extension of temporal ISC to multi-voxel pattern analysis (Haxby et al., 2014; Norman et al., 2006). Generally, the ISC is implemented with either a leave-one-out framework, in which one subject’s time course is correlated with the average of all other subjects for each region, or a pairwise framework, in which correlation analysis is performed between each possible pair of subjects (Finn et al., 2020). A limitation of this computational procedure is that the resulting correlations are highly interdependent and violate the assumption of common parametric tests (Nastase et al., 2019), requiring careful attention to the inference method. In addition, multiple cognitive and affective processes emerge in brain response to complex naturalistic stimuli (Bartels and Zeki, 2004; Simony and Chang, 2020). ISC reflects similarity in how each brain region encodes the stimulus across participant, not how a brain region(s) may response to different features of the natural stimuli based on the current ISC model. In order to mitigate these limitations, we proposed a novel tensor component analysis (TCA) framework that characterizes shared spatio-temporal patterns as well as idiosyncratic loadings of evoked activity to naturalistic stimuli across subjects in a purely data-driven manner that assesses all brain networks simultaneously.

While a matrix is a 2-D array, multidimensional data with more than 2 dimensions are referred to as tensors. TCA is a fundamental model for tensor decomposition, also known as tensor Canonical Polyadic Decomposition (CPD) (Kolda and Bader, 2009), that has been shown to have superior performance in identifying hidden signal sources compared with matrix decomposition, e.g. principle component analysis (PCA) and independent component analysis (ICA), applied to multidimensional data (Williams et al., 2018). Assuming the data meet the assumption of a mixture model, in which signal sources undergo a linear mixing process, TCA can more accurately estimate sources than matrix decomposition algorithms, and without any constraints. FMRI signals are innately multidimensional property and can be naturally represented in tensor form. In this study, a third-order fMRI tensor is organized as space × time × subjects (the order of modes does not impact the estimation). Through TCA analysis, the spatial and temporal information regarding brain network activity evoked by different stimuli exists in the first two dimensions. Subject loadings exist in the third dimension, which are used to investigate subject variability.

TCA has shown promise in a range of neuroscience researches. In the field of EEG signal analysis, nonnegative constraint TCA was applied on time-frequency domain data to identify event-related (Cong et al., 2015a, 2015b; Wang et al., 2018) and naturalistic stimulus-evoked EEG response (Zhu et al., 2020b). When TCA was applied for fMRI connectivity analysis, Mokhtari et al. (2019) explored the difference in the results of different tensor organizations and Zhu et al. (2020b) have explored the effectiveness of TCA for MEG data analysis. TCA has also been applied for fMRI data analysis (Andersen and Rayens, 2004) and has been adapted for multi-subject fMRI data analysis by placing spatial and temporal constraints to address inter-subject variability (Beckmann and Smith, 2005; Helwig and Hong, 2013; Kuang et al., 2020, 2015; Mørup et al., 2008; Zhou and Cichocki, 2012). We advance TCA for analysis of multi-subject fMRI data collected during naturalistic stimuli viewing by proposing a pipeline that does not place any constraints on the data.

In order to characterize stable shared spatio-temporal components across subjects with TCA, the reproducibility of TCA components is evaluated using a novel clustering method, tensor spectral clustering and model order (number of extracted components) is selected in terms of algorithm stability. The effectiveness of the proposed framework is demonstrated with simulated and traditional task fMRI in which we know the ground truth stimulation (and hence brain activity) timecourses. Then we apply the proposed framework to fMRI data collected during movie watching, in which there is no a *priori* model of brain activity, to identify spatial brain networks engaged during the task.

The rest of the paper is structured as follows. In section 2, Materials and Methods, we present the TCA model that was used to estimate spatio-temporal shared components, the decomposition algorithm used in this paper, the criteria that were used to evaluate the reproducibility of estimated components and the model order selection method, and the test datasets. In section 3, Results, we show the simulation and motor task fMRI results. After establishing the robustness of the proposed framework, we show results from applied our analytic approach to a naturalistic stimuli fMRI dataset from 184 participants, collected by the Human Connectome Project (HCP). In section 4 and 5, Discussion and Conclusions, respectively, we discuss our findings and the conclusions from our work. The basic principle of tensor spectral clustering that is used to evaluate the stability of estimated components is introduced in Appendix.

## Materials and Methods

### TCA model

The full strategy of TCA for naturalistic stimuli fMRI data consisted of three steps as shown in Fig. 1. Firstly, extract activity timecourses with independent component analysis (ICA) parcellation scheme; then, stack all subjects’ timecourses to construct a third-order fMRI tensor; finally, estimate spatio-temporal patterns and corresponding subject loadings with TCA.

**Figure 1.**
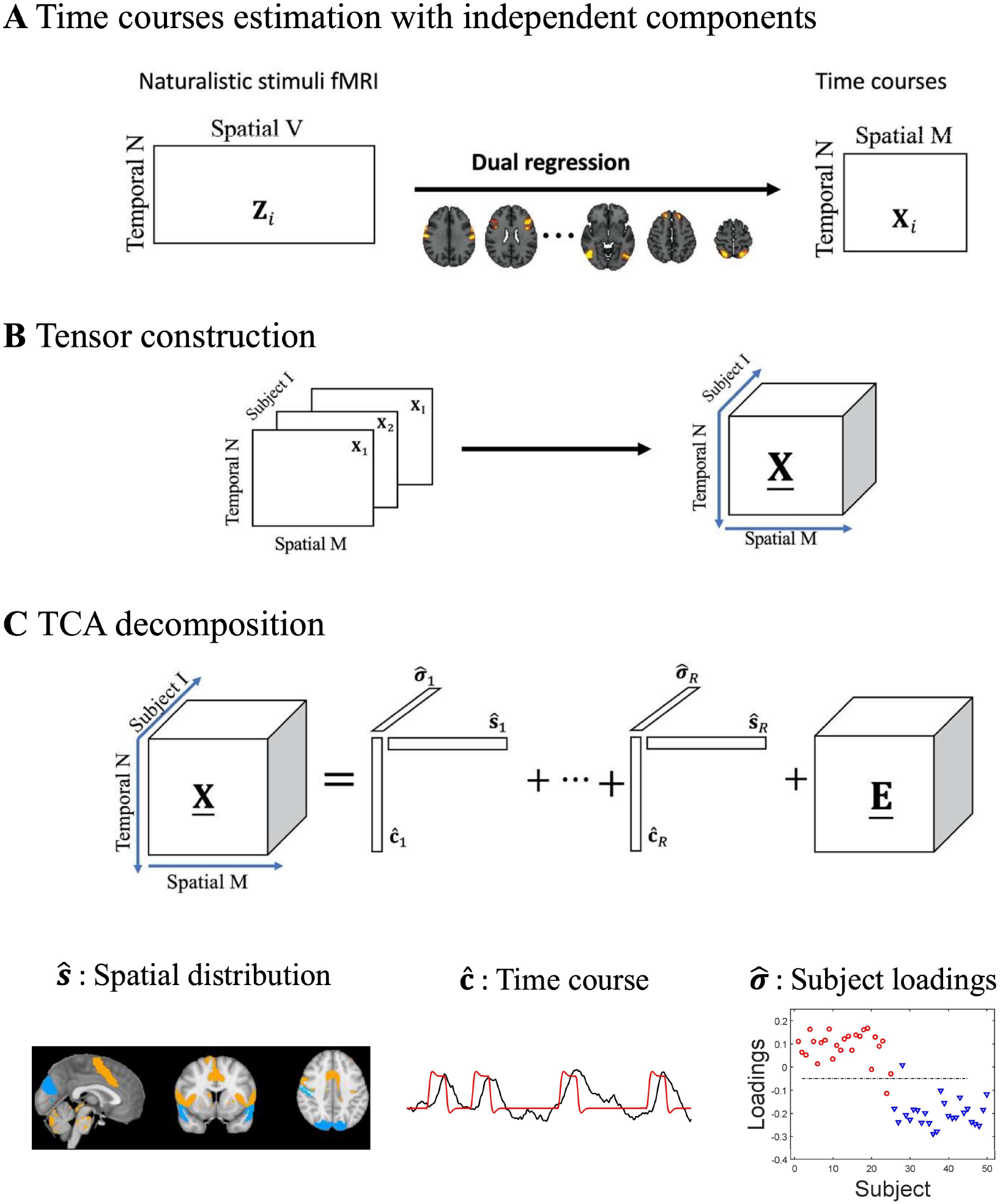
Tensor component analysis pipeline. (A) Estimating node timeseries with dual regression from standard preprocessed dense data. (B) Stacking all subjects’ node data to construct a tensor. (C) Extracting tensor components with tensor component analysis. The estimated components of each mode ŝ, ĉ, 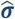 correspond to the spatial distribution, time course and subject loadings, respectively.

Rather than applying TCA to voxel-wise fMRI data, a whole brain parcellation scheme is applied to extract node (or region) timecourses for the TCA. This is done for several reasons. First, intensity of BOLD fMRI data is expected to be smooth across neighboring voxels and, with an appropriate parcellation method, node timecourses will have a higher signal noise ratio (SNR) compared with the SNR of timecourses from dense data (Glasser et al., 2016). Using ICA with high model orders (>100 to several hundred) will result in components that feature individual small brain regions, bilateral brain regions, or sparse sub-networks and can thus be considered as nodes for use in network analysis (Smith, 2012). Several studies have demonstrated that time course of spatial independent components can identify intrinsic brain networks (Allen et al., 2014; Jafri et al., 2008; Smith et al., 2012). Other work has also shown that brain parcellation with spatial ICA demonstrates better performance for network modeling compared with other parcellation methods (e.g., anatomical parcellations) and higher model order leads to better performance (Pervaiz et al., 2020). In this study, a brain parcellation scheme derived from ICA of the Human Connectome Project (HCP) resting state fMRI data with model order of 300 (provided by the HCP, Van Essen et al., (2013)) is used to extract timecourses from nodes for TCA. The timecourses are extracted via the first stage of dual regression (Nickerson et al., 2017) in which the full set of ICA components, **S**_ICA_, are regressed against each participant’s 4D fMRI data (e.g., multivariate spatial regression) to extract the timecourses. This is different from how conventional binary parcellation masks are used to extract average timecourses from each node that are then fed into the TCA in that the multivariate spatial regression accurately handles any potential spatial overlap among the ICA maps due to individual brain regions participating in more than one brain network (or sub-network) represented in the ICA maps.

#### Multilinear mixing model

For naturalistic stimuli fMRI we assume, similar to ISC model (Finn et al., 2020; Nastase et al., 2019), that time courses **C** ∈ ℝ^N×J^ (N is the number of time points, J is the number of patterns) and spatial patterns **S** ∈ ℝ^M×J^ (M is the number of nodes, 300 in this study) of nodes in the parcellation are stimulus-evoked responses that are consistent across subject. However, in contrast to the ISC model, there may also be J patterns that are shared across subjects that correspond to evoked responses associated with different features of the naturalistic stimuli. Thus, in our TCA-based model, for each pattern *j*, the loading ***σ***_*i,j*_ for subject *i* is different from other subjects. In this case, for subject i, the node timecourses **X**_*i*_ ∈ ℝ^M×N^, can be represented as:

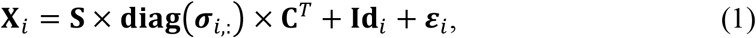

where **Id**_*i*_ is the stimulus-evoked response that is idiosyncratic to each subject. ***ε***_*i*_ corresponds to the spontaneous and noise component, which may reflect spontaneous neural activity and noise from non-neural physiological and scanner sources. **diag**(***σ***_*i*,:_) is a square matrix of order J with the elements of ***σ***_*i*,:_ on the diagonal and the other elements of the matrix is zero.

#### Tensor construction

A tensor is constructed from the multi-subject data, **<u>X</u>** ∈ ℝ^M×I×N^, which is the data from each subject's **X**_*i*_ stacked in the subject dimension. **I** is the total number of subjects. Although the spatial and the temporal patterns are assumed to be shared across participant, the loadings of the pattern in each participant are different. Of note, the model also does not place any assumption on the distribution of temporal courses or spatial distributions.

#### TCA unmixing model

TCA, also known as CPD, is a basic model for tensor decomposition. With different constraints, different algorithms can be derived, including non-negative canonical polyadic decomposition (NCPD) (Zhou et al., 2014) and independent constrained CPD (e.g. tensor-ICA Beckmann and Smith, 2005). In the TCA model, a third-order tensor **<u>X</u>** can be represented as the sum of several rank-1 tensors and a residual tensor **<u>E</u>** (Hitchcock, 1927), which is illustrated in Fig.1 (C). The mathematical formula is as follow:

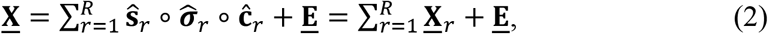

where vectors **ŝ**_*r*_ ∈ ℝ^M×1^,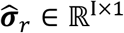 and **ĉ**_*r*_ ∈ ℝ^N×1^ are *r*^th^ estimated tensor component (TC) spatial distribution **S**, subject loadings ***σ*** and temporal courses **C** respectively. The operator ∘ represents the outer product of vectors. **<u>X</u>**_*r*_ represents the rank-1 tensor that constructed by the corresponding components of each mode. The idiosyncratic stimulus-evoked responses, spontaneous and noise components are contained in the residual tensor **<u>E</u>** and *R* represents the number of extracted patterns. Ideally, the number of tensor components, or model order, should be equivalent to the number of patterns, J, shared across subjects. However, in real-world applications, the number of patterns in the brain is unknown. In this study, model order is determined according to algorithm stability. For each tensor component, the voxel-level spatial distribution can be back reconstructed as **S**_ICA_ × **ŝ**_*r*_, the time course is ĉ_*r*_, which is the same across subjects. Differences of subjects exist at 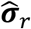.

### TCA estimation algorithm

The alternating least-squares (ALS) algorithm (Cichocki et al., 2015; Kolda and Bader, 2009) is used to estimate factor matrices **S**, **C** and ***σ***. In ALS algorithm, two of the factor matrices are fixed to optimize over the third factor matrix. For example, while time courses **C** is being estimated, the spatial distribution **S** and subject loadings ***σ*** should be fixed. The spatial distribution can be updated with the following rules:

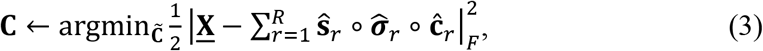

where *F* represents the Frobenius norm. The updating rule can be solved as a linear least-square problem that is convex and has a closed-form solution. The other factor matrices can be solved with the same updating rule. Three factor matrices were alternating updated until one of the stop criteria is met. The stop criteria can be defined as the absolute different value of data fitting of the adjacent two iterations was less than 1e-8 or the maximum number of iterations was more than 1000. The ALS algorithm can be free accessed from tensor toolbox (https://www.tensortoolbox.org).

### Algorithm stability analysis

Improving the reproducibility of neuroscience research is one of great concern (Poldrack, 2019; Poldrack and Farah, 2015). In this study, in order to assess the reproducibility of estimated rank-1 tensor 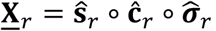, a novel clustering method, tensor spectral clustering, was proposed. In the stability analysis of TCA algorithms, for the given dataset, the same algorithm with the same parameters will be run *K* times with different initial conditions. When the number of extracted components is selected as *R*, for each mode *R* × *K* components can be estimated. Then the similarity matrices across these components of each mode **W**^(**S**)^, **W**^(**σ**)^, **W**^(**C**)^ ∈ ℝ^RK×RK^ would be fed into tensor spectral clustering, which is a co-clustering method that can fuse and assess the stability information of different modes simultaneously. Details of formulations of tensor spectral clustering can be found in the Appendix. In tensor spectral clustering, the number of clusters is defined as exactly same with the number of extracted components *R*. The stable component would produce a tight cluster. The stability index is quantified with the average intra-cluster similarities. Ideally, if the estimation of the component is stable, the inner similarity of the corresponding cluster is close to 1. The stability index of unstable components is approach to 0. The algorithm stability is defined as the average of components stability indices.

### Model order selection

Same as ICA, model order (number of extracted components) selection is a significant methodological concern when data driven algorithm is applied for fMRI data analysis (Abou-Elseoud et al., 2010; Beckmann, 2012; Kuang et al., 2018). When the selected model order is appropriate to the tensor to be decomposed, the algorithm would also be stable. In this study, we performed the tensor decomposition algorithm use a range of model orders. The algorithm stability under each model order was evaluated. The components under the model order with the highest algorithm stability index were used for further analysis.

### Simulations

In this section, we showcase the effectiveness of the proposed framework using numerical experiments which is performed in MATLAB. The spatial distribution of 29 ICA components (SimBT (Erhardt et al., 2012) http://mialab.mrn.org/software) and time courses **C** ∈ ℝ^100×8^ (http://mlsp.umbc.edu/simulated_fmri_data.html) are shown in Fig. 2. Four timecourses, as shown in the first row of Fig.2, represent consistent components across subjects. There are also three idiosyncratic components for each subject, which denote the scanner or motion noise consisted in fMRI scanning, as shown in the second row of Fig 2. These components would randomly circle shift for each subject. The timecourse in the third row of Fig 2 represents spontaneous brain activity included in the simulation. The total signal noise ratio (SNR) is controlled to be 2dB. We randomly generated node weight matrix **S** ∈ ℝ^29×8^ and subject loading matrix ***σ*** ∈ ℝ^10×8^ to generate 10 participants’ data. For each subject, the data can be created with equation (1). Then stack all subject data to construct a three-order tensor **<u>X</u>**_Simulation_ ∈ ℝ^29×10×100^. Then the tensor was decomposed with the model order range from 2 to 10. Under each model order, the TCA algorithm ran 20 times to evaluate the stability of the algorithm. The stability index is calculated with tensor spectral clustering. The correlation coefficient between estimate component and the corresponding ground truth works as a criterion to evaluate the performance of the proposed framework.

**Figure 2.**
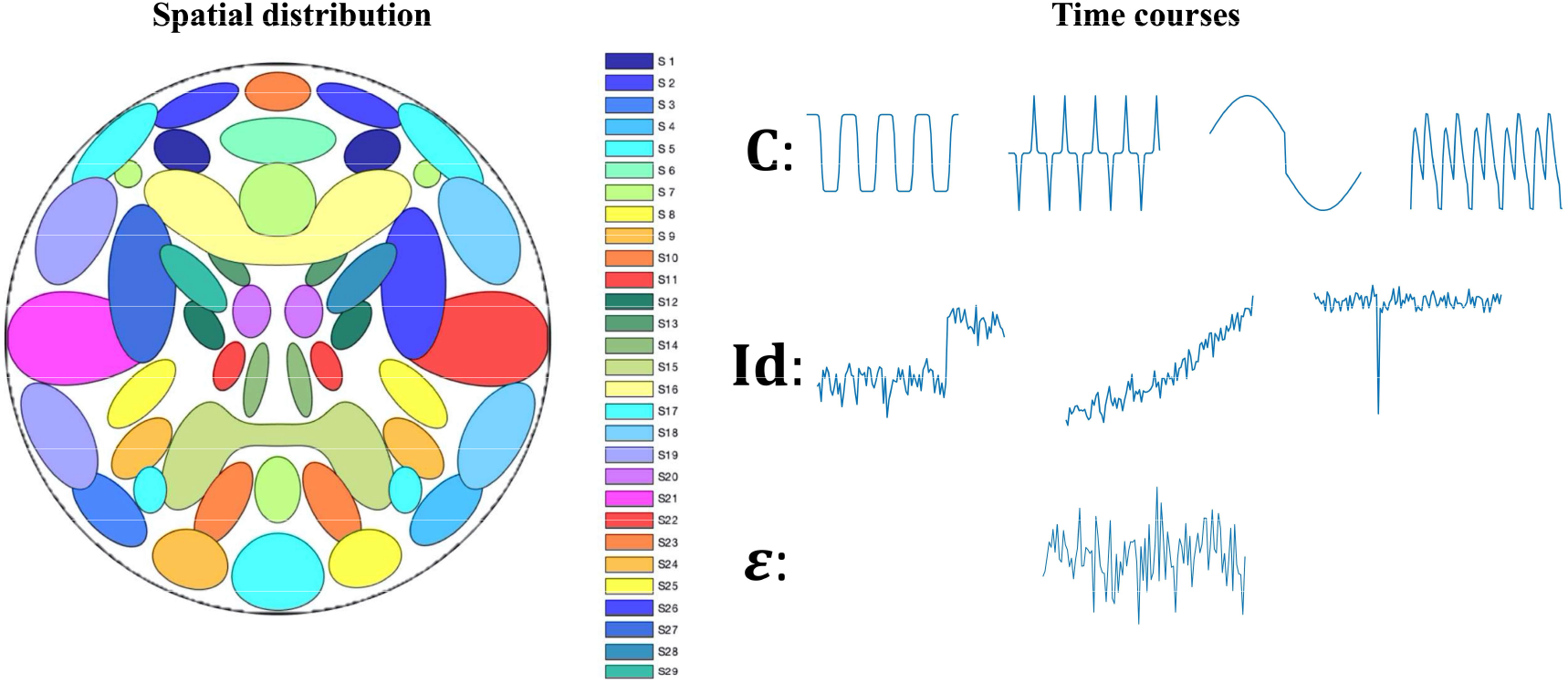
Ground truth spatio-temporal sources signal used in simulation. The left figure shows the spatial distribution of 29 ICA components. The right column shows the time courses to mimic the consistent components across subjects (the first row, **C**), the idiosyncratic components (the second row, **Id**) and the spontaneous components (the third row, ε).

### Motor task fMRI experiment

The effectiveness of the proposed framework is also demonstrated with traditional motor task fMRI in which we know the ground truth stimulation (and hence brain activity) timecourses. Randomly selected 100 healthy unrelated subjects (22-36 years) were utilized from the WU-Minn Human Connectome Project (HCP; Van Essen et al., 2013). During the motor task, participants are presented with visual cues that ask them to either tap their left or right fingers, or squeeze their left or right toes, or move their tongue to map motor areas. Each block movement type last 12 seconds and is preceded by a 3 second cue. For each subject total 284 scans were collected with TR=0.72s in a 3T scanner. Details of motor task of HCP data could be find in Barch et al. (2013).

The collected fMRI data went through the standard preprocessing pipeline (motion correction, distortion correction, highpass filtering, and nonlinear alignment to MNI template space (Glasser et al., 2013)) to the standard brain space. Time courses of visual cues for each type of movement were convolved with Hemodynamic Response Function (HRF) and went through a low-pass filtering with a high-frequency cutoff of 0.1Hz. The convolved time courses, as shown red lines in Fig. 4, are used to identify components extracted with TCA. Following the recommendations of previous study (Pervaiz et al., 2020), spatial ICA components were used for brain parcellation. Spatial ICA components are provided by the HCP under number of components of 300 and subject-specific time series for each node were derived using dual regression (Nickerson et al., 2017), as shown in Fig. 1 (A). Then time courses were detrended linear, quadratic, and cubic trends, and low-pass filtered with a high-frequency cutoff of 0.1Hz.

The data for each subject was then stacked on subject dimension to create a tensor **<u>X</u>**_Motor_ ∈ ℝ^300×100×284^ to be decomposed. Then the tensor is decomposed with TCA under model order from 2 to 20. And the same decomposition under each model order would be run 20 time to make sure the reproducibility of the estimated components. Then the model order is selected with the proposed model order selection method based on algorithm stability analysis strategy.

### Naturalistic stimuli fMRI experiment

The naturalistic fMRI dataset is also from the HCP (Van Essen et al., 2013). Subjects (healthy, 22-35 years) engaged in a movie-watching paradigm (voxel size = 1.6mm^3^, TR=1s) functional MR scanning in a 7T scanner. The sample used here (n = 184) reflects all available data for this paradigm. Each subject watched four 15-min movie runs (MOVIE1-MOVIE4). Each run comprised five video clips presented in a fixed order. The fMRI data collected during validation clip (the fifth video clip) was used in this study. The validation clip is consisted of a montage of brief (1min 22s) moving scenes depicting people and landscapes. The music that along with movie scene also contains plenty of information that may evoke consistent brain activities across participants (Alluri et al., 2012). Five music features (Fluctuation Centroid, Fluctuation Entropy, Key Clarity, Mode and Pulse Clarity) were extracted using the MIRToolbox (Lartillot, O., 2007). With the proposed framework, dynamic temporal alteration of brain networks can also be estimated. These music features facilitate the understanding and interpretation of the estimated tensor components. With the proposed framework, a component that related to social scene is identified. The subject loadings of the component were used to investigate the relationship between behavior data Antisocial Personality Problems Raw Score and brain activities. The behavior data is a score that evaluates the antisocial personality disorder assessed using the Semi-Structured Assessment for the Genetics of Alcoholism (SSAGA; BUCHOLZ et al., 1994; Hesselbrock et al., 1999).

All fMRI analyses began with the FIX-denoised data, which includes standard preprocessing (motion correction, distortion correction, high pass filtering, and nonlinear alignment to MNI template space (Glasser et al., 2013)) plus regression of 24 framewise motion estimates (six rigid-body motion parameters and their derivatives and the squares of those 12) and regression of confound components identified via independent components analysis (Griffanti et al., 2014; Salimi-Khorshidi et al., 2014). Details of data acquisition and basic data preprocessing can be found in previous studies (Glasser et al., 2013; T. Vu et al., 2016; Van Essen et al., 2012). The tensor organization procedure is exactly same with that of motor task fMRI, as shown in Fig. 1 and the tensor to be decomposed is **<u>X</u>**_Naturalistic_ ∈ ℝ^300×184×82^.

In this experiment, our goals are three-fold. Firstly, apply the proposed framework to all subjects to explore what kinds of brain activity can be evoked with the movie stimuli. All subjects’ fMRI data went through the proposed framework. Then the relationship between the estimated components and subjects’ behavior data as well as music features was further analyzed. Secondly, to investigate the reproducibility of the estimated components, the subjects were randomly and equally divided into two cohorts. And apply the proposed framework on these two cohorts separately. Then evaluate the consistency of the estimated components from these two cohorts. Thirdly, two version of movie files compiled as stimuli were used in MOVIE4. In the posterior movie stimuli, 4 frames were added before the validation clip, the deviation of two version movie is only 167ms, no more than 1 TR. To evaluate the performance of the proposed framework on exploring subjects’ difference, 25 subjects of each type of movie stimuli were randomly selected. Then these 50 subjects’ fMRI data went through the proposed framework, to probe whether the conditions’ difference can be detected in subject loadings. And performance of TCA and ISC are also compared in this experiment in terms of conditions’ difference detection. In ISC, leave-one out method, in which one subject’s time course is correlated with the average of all other subjects for each region, was applied to calculate the subjects’ score.

## Results

### Simulations

Fig. 3 shows the simulation results. Fig. 3(a) exhibits algorithm stability indices under different model orders. We could find that the algorithm stability curve reaches its high at the model order of both 3 and 4. Since the model order selected as 4 means one more component can be extracted, the tensor components at the model order of 4, which is exactly same with the number of consistent components across subjects, were further analyzed. Note that the algorithm stability index at the model order is 1, which means all components estimated on the model order from different runs are the same and all these components are repeatable. The spatial distribution, time courses and subject loadings of each component are exhibited in Fig.3 (b-e). Based on the correlation coefficient between estimated components and ground truths, we could find that all four components are successfully estimated. In order to highlight the activated and deactivated brain regions, the spatial distribution of the estimated component was exhibited with proportional threshold (10%, Garrison et al. (2015)). In spatial distribution, warm colors represent activation (relative to global average) while cool colors represent deactivation within each component.

**Figure 3.**
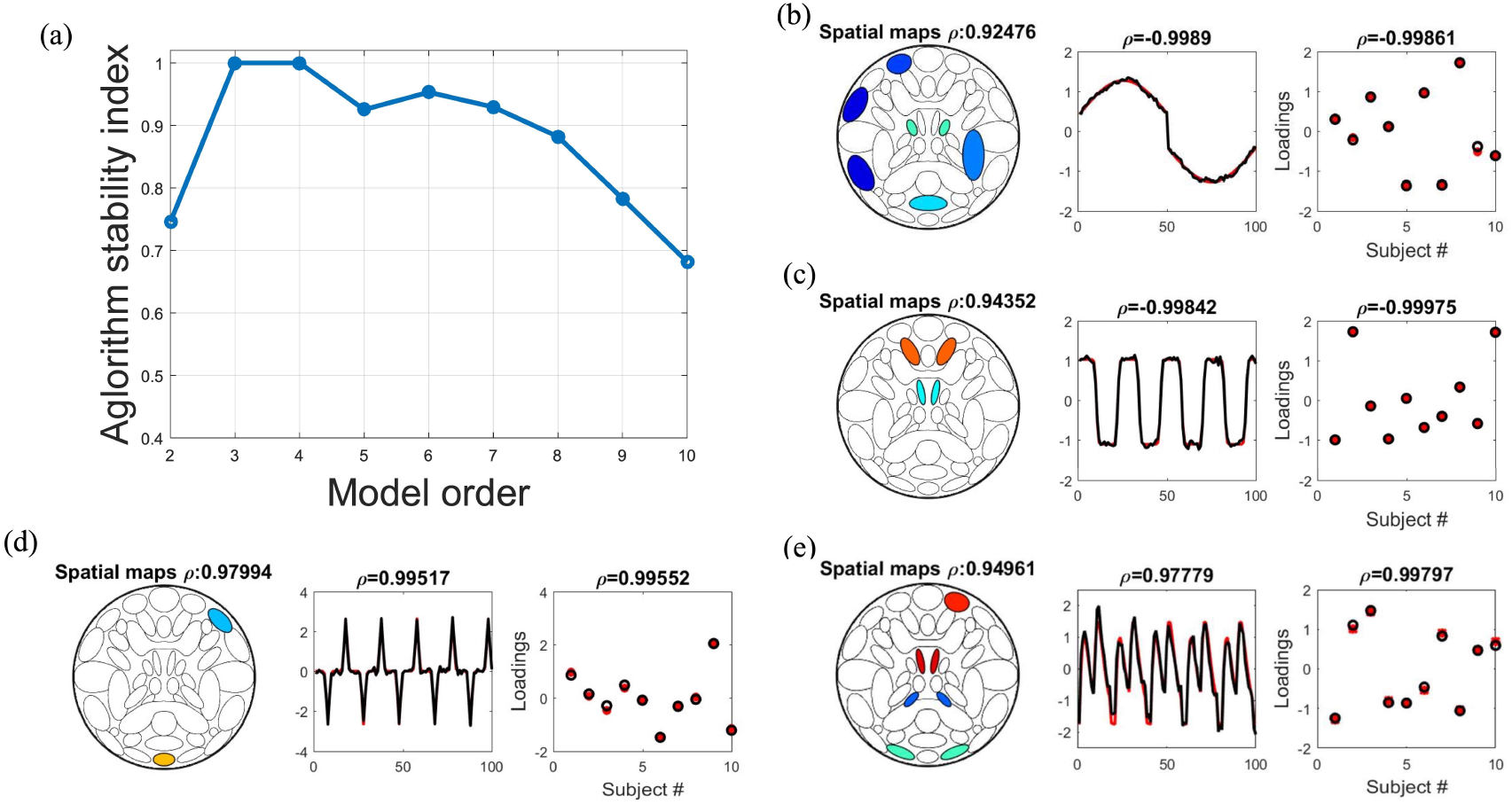
Simulation results. (a) Algorithm stability under different model orders. The algorithm reaches the highest stability at the model order of 4, which is exactly same with the number of consistent components. (b) - (e) estimated components and ground truths. The symbol ρ means the correlation coefficient between estimate component and the corresponding ground truth. In spatial distribution, proportional threshold (10%, Garrison et al. (2015)) was used and warm colors represent activation (relative to components global average) while cool colors represent deactivation (relative to components global average) within each component.

### Motor task fMRI experiment

For motor task fMRI data, with the proposed framework, the algorithm stability curve reaches its peak at the model order of 5. Hence, the estimated components at the model order of 5 were further analysis.

The spatial distribution, time courses of the estimated components are demonstrated in Fig. 4. The spatial distribution was also proportional thresholded (10%, Garrison et al. (2015)) and the warm colors represent activation and the cool colors represent deactivation. For each component, after standardization, both stimulus timing and estimated time courses are demonstrated in the same subfigure to identify of the estimated components. In the experiment, the embedded components that correspond to the tongue (TC#1), foot (TC#2) and total movement (TC#3) are successfully estimated. However, the components that corresponding to the hand movement is failed to be detected. For the tensor component that corresponding to the tongue movement (TC#1), the brainstem plays an important role. The postcentral gyrus dominates the movement of foot (TC#2). For total movement tensor component, sensorimotor area is activated (TC#3).

**Figure 4.**
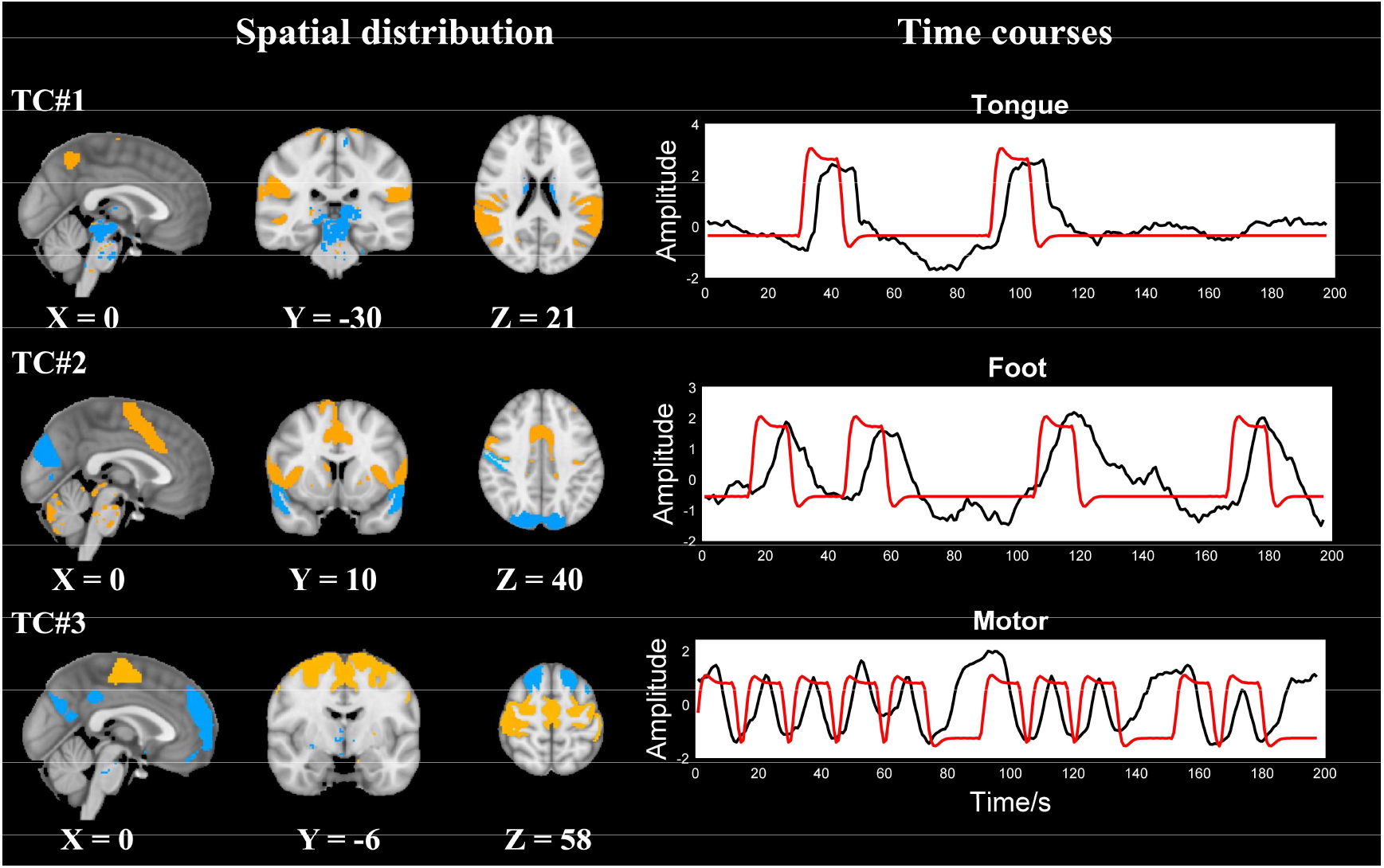
The spatial distribution (left) and the time courses (right) of estimated components from motor task fMRI (TC: tensor component). In spatial distribution, warm colors represent activation (relative to within-state global average) while cool colors represent deactivation (relative to within-state global average) within each state. In time courses, the black line represents estimated time courses, and the red line represents visual cues after being convolved with hemodynamic response function.

Fig. 5 shows the estimated components with similar time course but different spatial distribution. It worth note that, even though the time courses are similar between TC#3 and TC#4, different spatial distribution split them into different components. Compared with TC#4, sensorimotor area is activated in TC#3, while visual area plays more important roles in TC#4. We believe that TC#3 is related to motor decision making and the TC#4 is evoked with stimulus cues.

**Figure 5.**
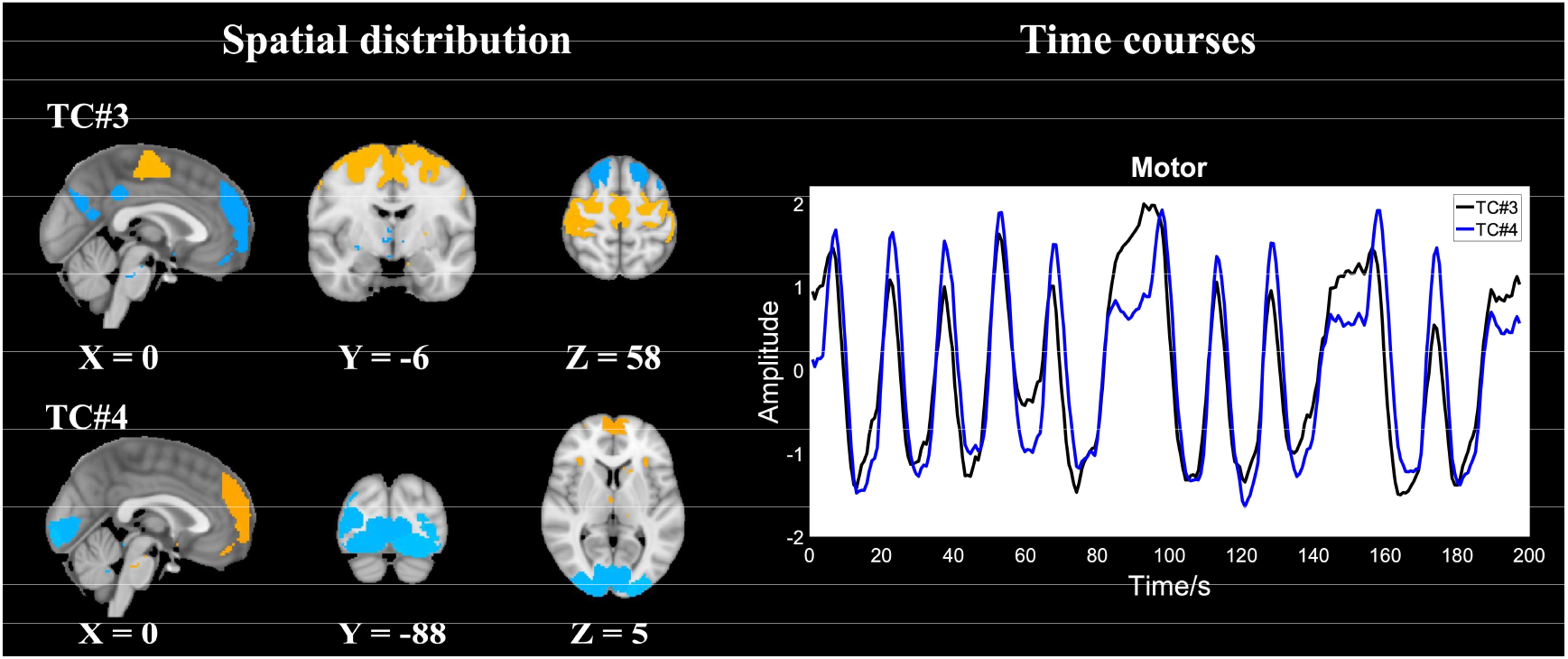
The estimated components with similar time course but different spatial distribution. The TC#3 corresponding to the motor decision making and the TC#4 represent the brain activity evoked with visual cues.

### Naturalistic stimuli fMRI experiment

For naturalistic stimuli fMRI, the TCA algorithm reaches the highest stability at the model order of 3. The estimated components under the model order are further analyzed.

The spatial distribution, time courses and subject loadings of the first tensor component are shown in Fig. 6 (a). Fig. 6 (b) shows the ISC estimated spatial pattern with the same data. Bilateral occipital fusiform gyrus, lingual gyrus and superior temporal gyrus are covered by both TCA component and ISC result. However, bilateral postcentral gyrus and superior parietal lobule and left precentral gyrus are also found in ISC result but not in TCA component. Bilateral cerebellum and lateral occipital cortex only exist in TCA component. Fig. 6 (c) shows the correlation coefficient of subject temporal courses of lingual gyrus and temporal gyrus, which are both covered by TCA component and ISC result, with time course of the first tensor component estimated with tensor component analysis. The bar plot of Fig. 6(c) shows that the time course of lingual gyrus is negative correlated with the estimated time course, but the time course of temporal gyrus is positive correlated with the estimated time course, which demonstrates that under the movie stimuli lingual gyrus and temporal gyrus with opposite activation pattern. TCA component successfully demonstrate it but ISC fails to.

**Figure 6.**
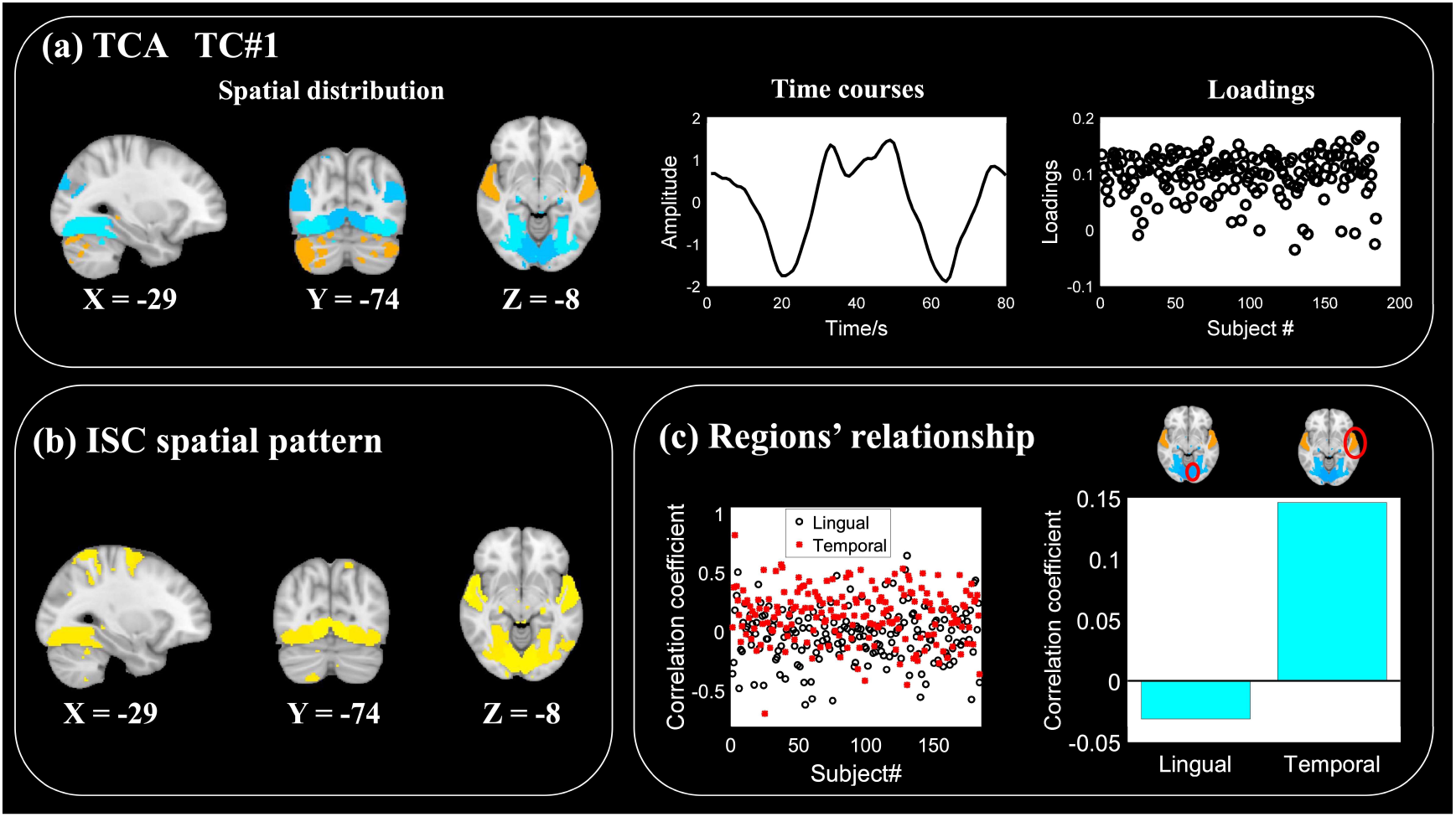
(a) The spatial distribution, the time course, and the subject loadings of the first estimated tensor components of naturalistic stimuli fMRI (TC: tensor component). (b) Spatial pattern estimated with ISC. (c) The scatter plot (left) shows the correlation coefficient of subject temporal courses of lingual gyrus and temporal gyrus with time course estimated with tensor component analysis (TCA). The bar graph (right) shows the average value of correlation coefficient.

For the TC#1, the estimated time course and the onset point in the stimuli movie is matched to show the relationship of them, as shown in Fig. 7 (a). We have a preliminary feeling that, the landscape scene will lead the curve goes up while the social related scene will make the curve goes down. Hence, we test the correlation coefficient between the subject loadings of this component and the social related behavior data. We found that it is significant correlated with score of antisocial personality problem as shown in Fig. 7 (b).

**Figure 7.**
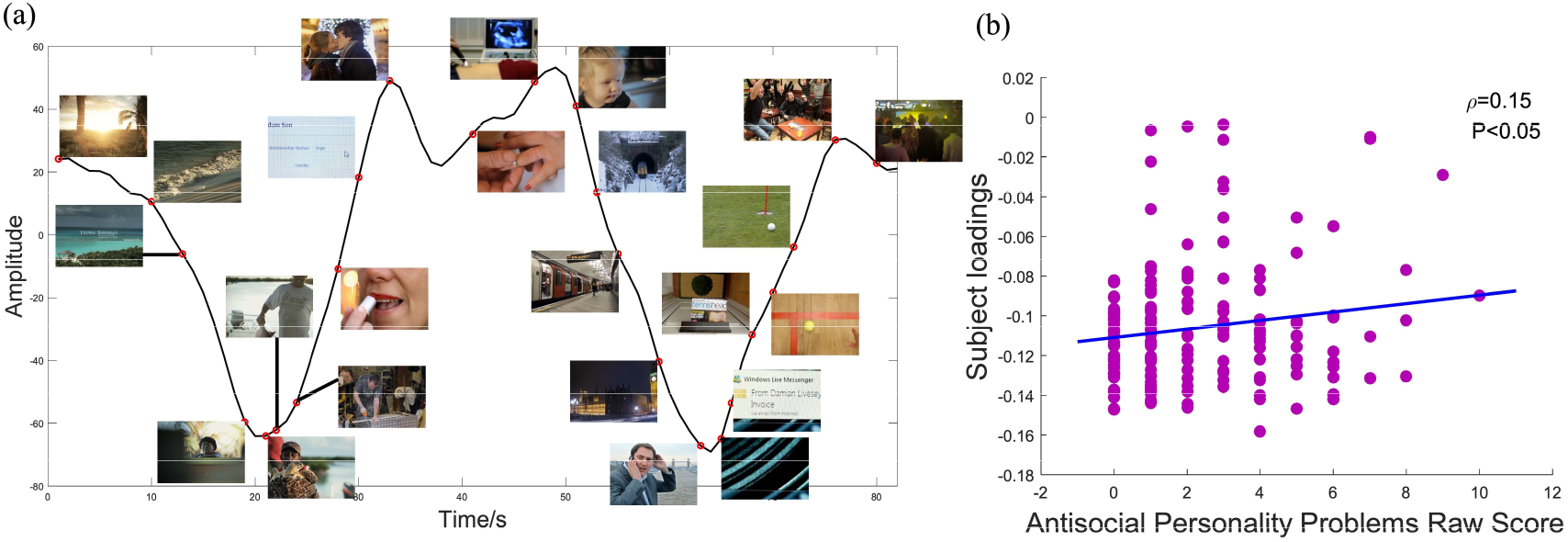
(a) Timeline of the first tensor component and the onset point of each scene. The trends of the timeline with landscape and social related scene are different. (b) The subject loadings of the first tensor component significant (uncorrected) correlated with Antisocial Personality Problems Raw Score (ρ: correlation coefficient).

Figure 8 shows the second and the third components estimated with tensor component analysis. The red lines in temporal subfigure (medium) show the time course of music features that significantly correlated with estimated time course. The time course of the second tensor component significantly correlated with the music feature Pulse Clarity and the time course of the third tensor component significantly correlated with the music feature Key Clarity. For all three estimated tensor components, bilateral cerebellum, occipital fusiform gyrus, and lateral occipital cortex are involved in. The lingual gyrus is deactivated in the TC#1 but not in TC#2 and TC#3. Different from the TC#1, left frontal pole is deactivated in both TC#2 and TC#3. The bilateral superior temporal gyrus exists in TC#1 and TC#2 but not in TC#3.

**Figure 8.**
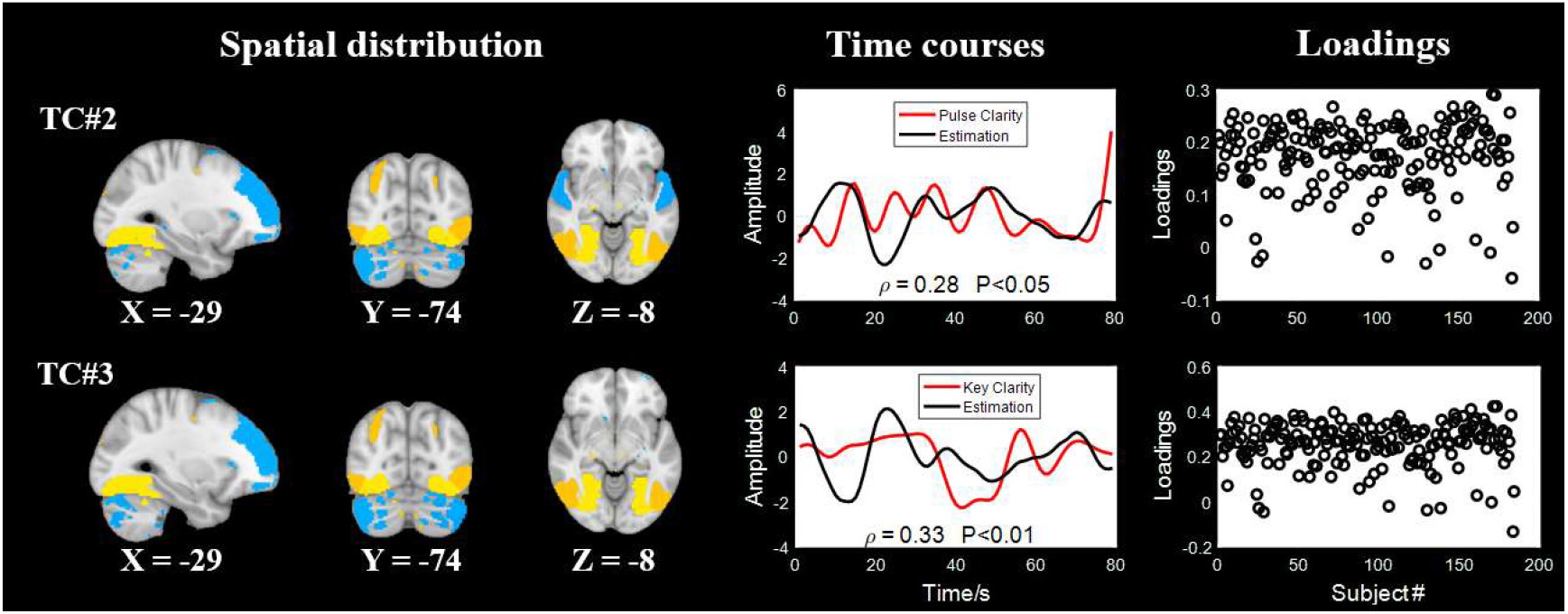
The second and the third components estimated with tensor component analysis. The red lines in temporal subfigure (medium) show the time course of music features that correlated with estimated time course significantly (ρ: correlation coefficient).

To test the reproducibility of the estimated component, the subjects were equally divided into two groups. These two groups went through the same TCA framework separately. As shown in Fig. 9, for both group, algorithm reaches the highest algorithm stability at the model order of 3. The right column of Fig. 9 demonstrates the consistency between two decomposition runs. It shows that the TC#1 and TC#3 from both cohorts are highly consistent, which means these components are evoked by the stimulus movie. We also observed that the consistency of the corresponding spatial distribution and find that correlation coefficients are also high (not shown in the paper). However, the consistency of the TC#2 of two cohorts is a little bit low.

**Figure 9.**
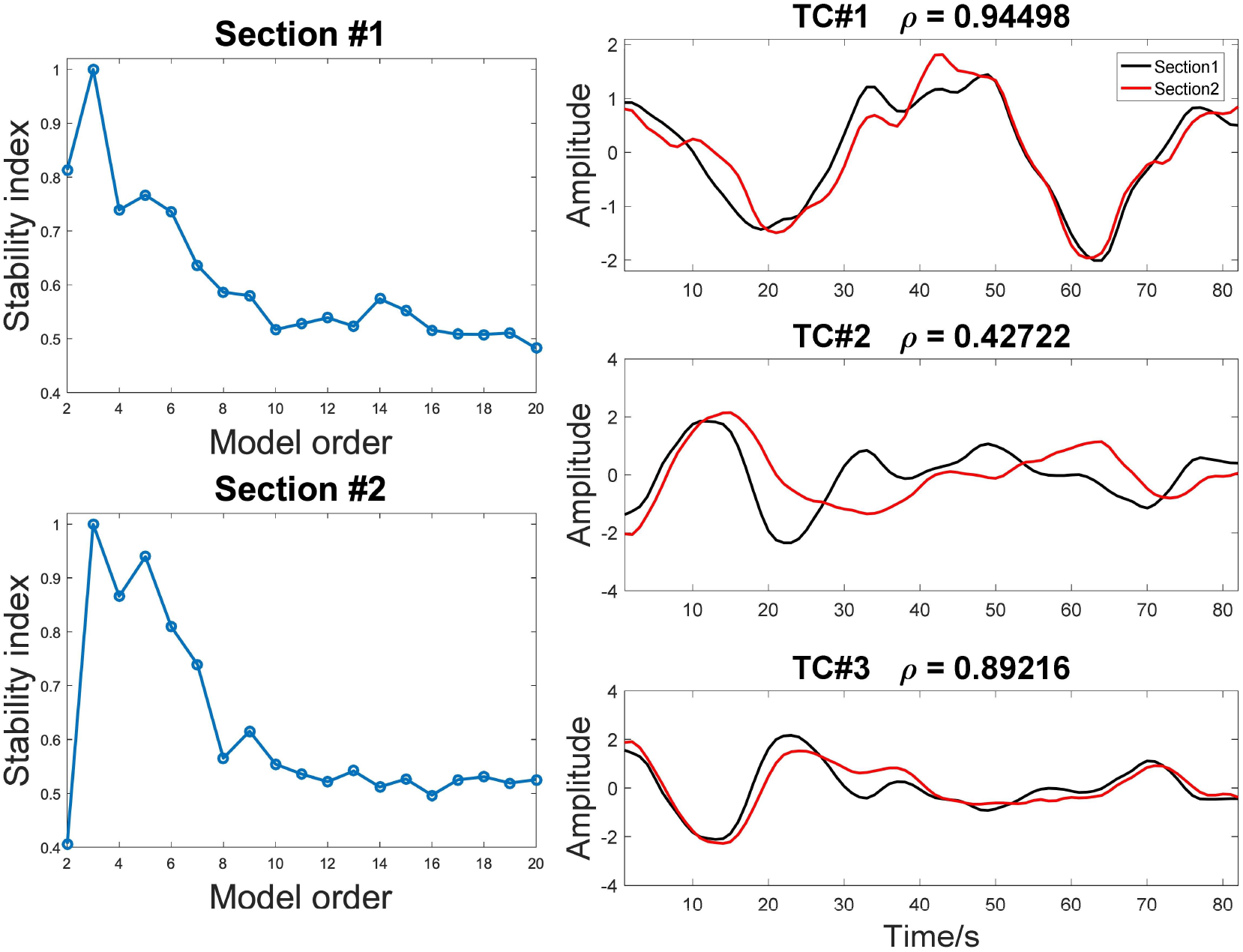
Reproducibility of the estimated components. The section#1 and section#2 represent different cohort of randomly divided subjects (ρ: correlation coefficient).

We also evaluated the power of the proposed framework in terms of subjects’ difference detection. The framework was applied on subjects that went through two with millisecond deviation movie stimuli and view it as two conditions. The results shown that subjects with different conditions are successfully distinguished with the estimated subject loadings, as shown in Fig. 10(c). The subjects at different conditions can be easily distinguished with a threshold value (dash line of Fig. 10(c)). In contrast, even though ISC can also identify spatial distribution (Fig. 10 (b)), which is similar with that of TCA component (Fig. 10(a)), conditions’ difference is destroyed by subject correlation (Fig. 10(d)).

**Figure 10.**
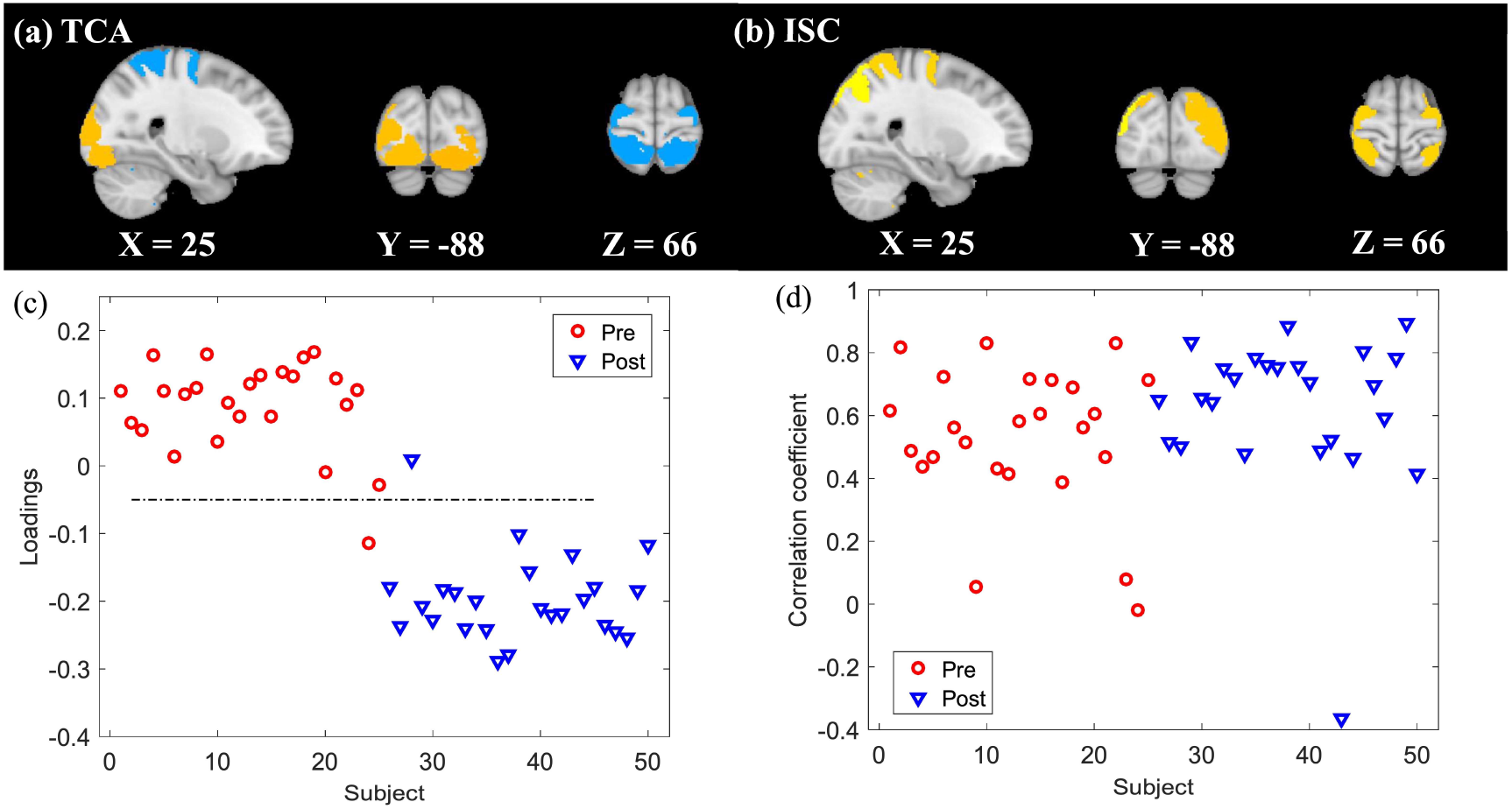
Comparison of TCA and ISC in terms of detection of individual difference. (a) Spatial distribution of the first component estimated with TCA. (b) Spatial distribution of ISC estimated consistent map. (c) Subject loadings of the first TC exhibit difference of experiment conditions (TC: tensor component). (d) Subjects’ score of ISC calculated with leave-one out method. The red circles represent loadings of subject of the first version movie stimuli. The blue triangles represent loadings of subject of the second version movie stimuli.

## Discussion

In this study, an analysis framework to discover shared spatio-temporal components across subjects from naturalistic stimuli fMRI with TCA was proposed. In the framework, a third-order tensor is constructed from the timeseries extracted from all brain regions from a given parcellation, for all participants, with modes of the tensor corresponding to spatial distribution, time series and participants. TCA will then reveal spatially and temporally shared components, i.e., naturalistic stimuli evoked networks, their temporal courses of activity and subject loadings of each component. The stability of the extracted components is evaluated with a novel clustering method, tensor spectral clustering, to guarantee the reproducibility of the results. Based on the algorithm stability under a range of model orders, the most suitable model order that makes algorithm stable can be recommended. Extensive experiments demonstrate that the framework is feasible, and the interpreted and reproduced components can be extracted.

Compared with traditional fMRI study designs, naturalistic stimuli are complex and dynamic, and it is much more difficult to generate a model of evoked activity for analyses. Under the movie stimuli, the timecourses of brain activity across subjects are consistent, which is changing along with movie plots going on. Besides, the specific shared brain network(s) would also be activated for the same kind of information processing. Based on this assumption, a TCA framework that can characterize shared spatio-temporal patterns of evoked activity to naturalistic stimuli across subjects was proposed. FMRI signals have an innate multidimensional property and can be naturally represented in tensor form. Then the tensor can be decomposed with TCA. The estimated components of each mode represent the spatial distribution, time courses and subject loadings, which exhibit participant difference. This work demonstrates that TCA can meet the naturalistic stimuli fMRI’s analysis needs and under the proposed framework, the brain activities evoked with naturalistic stimulus were extracted and the patterns that corresponding to different stimuli features is also distinguished (Fig. 6 and Fig. 8).

The proposed framework exhibits several merits. Firstly, different from ISC, the proposed framework can estimate subject loadings directly, which can be used to explore participants’ difference (Fig. 7(b)) or conditions’ difference (Fig. 10). Secondly, the specific brain activities response to features of stimuli can be separated (Fig. 6). Furthermore, the components that with the same time series but different spatial distribution can also be distinguished (Fig. 5). Thirdly, the time courses of estimated components can also be estimated, which facilitate the interpretation of the estimated components. Fourthly, combined with evaluation of algorithm stability, reproducibility of the estimation is promoted (Fig. 9).

With the proposed framework, TCA exhibits promising performance. For motor task fMRI, TCA successfully identify networks evoked by tongue, foot and total movement (Fig. 4). Even though the temporal courses are similar between visual stimuli and motor decision, TCA can successfully distinguish them. However, the algorithm fails to identify the components that evoked with hand movement. This may because that hand movement is sophisticated and the variance across subjects makes it difficult to estimate a common spatio-temporal pattern. This also reflects that the proposed framework can only be applied to extract the components that are highly consistent across subjects. More advanced method that can extract misalignment components across participants worth further investigation. Under the naturalistic stimulus, the various across subjects have attracted more and more attention (Finn et al., 2020; Nastase et al., 2019). With the proposed framework, individual difference can successfully be distinguished. Firstly, the component that related to the behavior data Antisocial Personality Problems Raw Score is identified with the proposed framework (Fig.7 (b)). Secondly, TCA successfully identify subjects with millisecond stimuli difference (Fig. 10 (c)). Compared with ISC, TCA can not only distinguish the participants’ difference but also the temporal courses of each pattern, which facilitate the interpretation of the estimated pattern. In terms of spatial distribution, TCA can not only estimate the naturalistic stimuli evoked spatial distribution but also the relationship of brain regions that are involved in the same brain network (Fig. 6 (a) and Fig. 6 (b)). This is because ISC only concern about the consistency across subjects but ignore the network spatial configuration.

Improving the reproducibility of neuroscience research is one of great concern (Poldrack, 2019; Poldrack and Farah, 2015). To guarantee the stability of the TCA algorithm, a novel spectral clustering algorithm was proposed and applied on the components estimated from multiple runs. In simulation experiment, the model order selected based on algorithm stability is exactly same with the number of consistent components across subject, which demonstrates that with the algorithm stability index appropriate model order can be recommended (Fig. 3(a)). In the naturalistic stimuli fMRI, the results demonstrate that subjects shared spatial-temporal components are reproduced with different individuals (Fig. 9). Actually, we also assessed the test-retest validity of the same subject went through the same naturalistic stimuli. However, the consistency is poor, which may because the neural processing alters with repetitive stimuli.

Naturalistic stimuli fMRI is a powerful tool to study brain network interactions during the daily life. Our results demonstrate that with appropriate movie stimuli and our proposed processing framework, the brain activity that significant correlated with Antisocial Personality Problems Raw Score is identified (Fig. 7). This shows the power of the naturalistic stimuli. The subjects with high antisocial score may fake their response when they go through the questionnaire or in abstract stimuli paradigm. However, the brain activity is hard to pretend in the naturalistic stimuli paradigm. Since the movie stimuli applied in this study is not designed for this purpose and the correlation coefficient value (Fig. 7(b)) will not significant after correction, a more suitable movie stimuli to evoke wanted components worth further investigation. Besides, only three shared spatial-temporal components can be estimated in the experiment. Even though naturalistic stimuli contain plenty of features, because of variance of subjects’ attention, there are not much brain activities shared across subjects. In the further study, the selection/designation of naturalistic stimuli need pay more attention to.

There are some limitations of this study. In this study, brain template comes from ICA with settled dimensionality. Several studies (Allen et al., 2014; Jafri et al., 2008; Pervaiz et al., 2020; Smith et al., 2012) have demonstrate that time course of spatial independent components can identify intrinsic brain networks. The performance of TCA with different templates (Glasser et al., 2016; Janes et al., 2019; Shen et al., 2013) was also compared and ICA template performs best. However, several studies have demonstrated that the model order can greatly impact on the estimated components (Abou-Elseoud et al., 2010; Beckmann, 2012; Kuang et al., 2018). We also tested the back-projection method to get the voxel level spatial distribution. Limited with selected model order, the ICA template is still not pure, which makes the voxel-level spatial distribution hard to be thresholded. Our previous study (Hu et al., 2020) has proposed an effective strategy, Snowball ICA, to address the dimensionality selection issue. In the further study, a more comprehensive template worth being further investigated with Snowball ICA. Even though the paper demonstrate that the proposed framework can distinguish subjects that went through millisecond deviation stimuli, how the framework can be applied on disease diagnosis, coordinate with appropriate naturalistic stimuli, still need further investigation.

## Conclusion

The proposed tensor analysis framework is a powerful method that can extract embedded components evoked with naturalistic stimuli. With the proposed framework, meaningful and reproduced tensor components can be extracted and subjects that with different conditions are successfully distinguished. Three experiments (simulation, motor task fMRI, naturalistic stimuli fMRI) demonstrated that the proposed framework is a promising tool to extract brain networks evoked with naturalistic stimuli.

## Acknowledgements

This work was supported by National Natural Science Foundation of China (Grant No.91748105), National Foundation in China (No. JCKY2019110B009 & 2020-JCJQ-JJ-252) and the Fundamental Research Funds for the Central Universities [DUT2019, DUT20LAB303] in Dalian University of Technology in China. This work was also supported by China Scholarship Council (No.201806060038).

## Appendix TCA stability analysis with tensor spectral clustering

## Background of spectral clustering on graph theory

In this study, a novel tensor spectral clustering algorithm is proposed to evaluate the stability of TCA algorithms. For ease of understanding, background of spectral clustering is demonstrated at here. Spectral clustering is a technique that roots from graph theory. Consider an undirected weighted graph G = (V, E). V is a set of nodes. E is a set of edges between nodes. The weighted adjacency matrix of G is denoted as a symmetric matrix **W**. The generalized degree of the vertices of G is defined as **D** = diag(**We**), where e is an all-ones vector. The combinatorial Laplacian matrix is defined as **K** = **D** − **W**. The transition matrix of the graph **P** = **W**^T^**D**^−1^ is a column stochastic matrix. Thus, the matrix could be interpreted as Markov chain. The stationary distribution of the Markov chain π = diag(**D**).

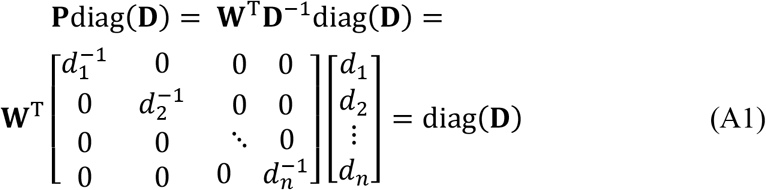

It also means that diag(**D**) is an eigenvector of **P** and the corresponding eigenvalue is one (R.Benson et al., 2015).

In the partition of a graph, it is assumed that the graph could be separated two parts S and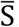. Bottleneck of a graph is the boundary of two clusters. The bottleneck ratio of the set Ψ is defined as:

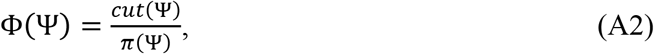

where 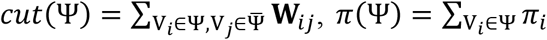. Small bottleneck ratio indicates a good partition of the graph (Levin et al., 2007).

The indicator vector f over the nodes in G is defined:

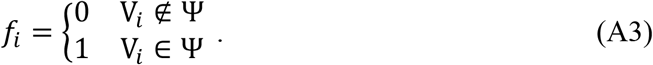

The property of Laplacian matrix is leveraged as follow:

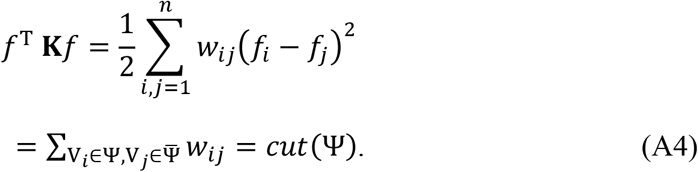

Since π = diag(**D**), *π*(Ψ) could be represented as *f*^T^ **D***f*. Hence, the objective of matrix spectral clustering is to find a *f* to minimize 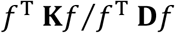.

Both matrices **K** and **D** are positive definite. According to generalized Rayleigh entropy, the solution is the vector *f* such that K*f* = λ**D***f*. We observed that:

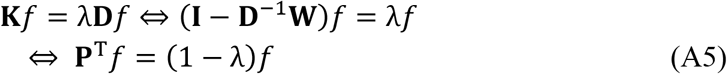

So, the problem becomes looking for the eigenvector of **P** (Ng et al., 2002).

## Tensor spectral clustering

For multimode tensor spectral clustering, different mode has different *transition matrix*. In this appendix, three modes tensor is used as an example for tensor spectral clustering. **P**^(1)^, **P**^(2)^, **P**^(3)^ are three transition matrices for three modes tensor decomposition. They are calculated as follow:

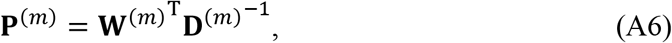

where **D**^(m)^ is defined exactly same with matrix spectral clustering. The generalized *transition tensor* **<u>P</u>** could be defined as:

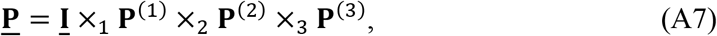

where **<u>I</u>** ∈ ℝ^RK×RK×RK×RK^ is a unit tensor, and the number of modes of **<u>I</u>** is one more than the number of modes of the tensor to be decomposed.

Same to matrix spectral clustering, generalized singular value decomposition (SVD), HOSVD (Lieven et al., 2000), is applied on *transition tensor* **<u>P</u>**. For HOSVD of **<u>P</u>**, the essence of the decomposition of the last mode is eigen value decomposition of covariance matrix of matrix unfolding of tensor **<u>P</u>** on the last mode. The unfolding of tensor **<u>P</u>** on the last mode is denoted with **<u>P</u>**_(end)_, which can be calculated as follow:

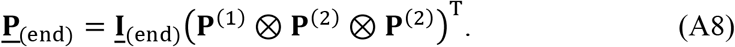

The covariance matrix of **<u>P</u>**_(end)_ is:

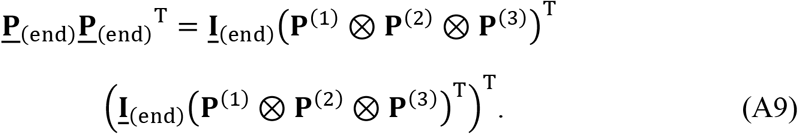

We defined that:

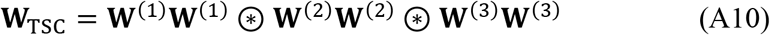

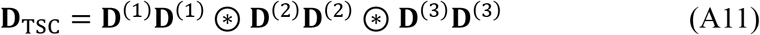

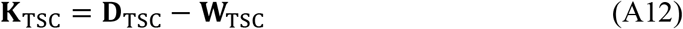

**W**_TSC_ is a symmetric matrix and represents weighted adjacency matrix of TSC. **D**_TSC_ is a diagonal matrix. Both matrices **K**_TSC_ and **D**_TSC_ are positive definite. Then the equation (A9) could be reduced as:

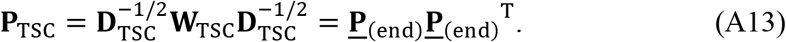

Same with matrix spectral clustering, the purpose of tensor spectral clustering is to find a *f* to minimize 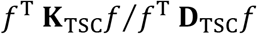. Based on the generalized Rayleigh entropy and diagonal property of **D**_TSC_, the objective function:

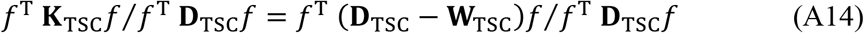

is equivalent to find a *f* to minimize 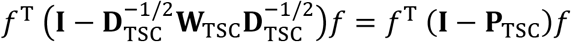

At this stage, the problem of tensor spectral clustering is reduced to matrix spectral clustering. So, the last mode eigenvector of HOSVD of *transition tensor* **<u>P</u>** could be used for multimode co-clustering.

Given a set of samples that we want to cluster into *k* subsets, each sample has more than one modality to be considered. The procedure of TSC is as follows:

1. Form the weighted adjacency matrices of each modality: **W**^(1)^, **W**^(2)^, **W**^(3)^, ⋯.
2. Define the *transition matrix* of each modality: **P**^(1)^, **P**^(2)^, **P**^(3)^, ⋯. Then the *transition tensor* **<u>P</u>** is defined with equation (A7).
3. Find the *k* eigenvectors ***v***_1_, ***v***_2_, ⋯, ***v***_*k*_ corresponding to the *k* largest eigenvalues of the last mode of transition tensor **<u>P</u>** with HOSVD. Form the matrix **V** = [***v***_1_, ***v***_2_, ⋯, ***v***_*k*_] by concatenating eigenvectors in columns.
4. Normalize each row of **V** to have unit length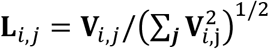.
5. Cluster points, each row of **L**, into *k* clusters via Hierarchical clustering (Gordon, 1987).
6. Find the corresponding row *i* of **L** and original sample, assign the original sample to the cluster *j* that row *i* assigned.

The TSC software is available at https://github.com/GHu-DUT/Tensor_Spectral_Clustering.

For the stability analysis of TCA algorithms, for the given dataset, the same algorithm with the same parameters will be run *K* times. Under the number of extracted components *R*, for each mode, there are *R* × *K* components. When TSC was applied in the stability analysis of TCA algorithms, the similarity matrices of each mode **W**^(**S**)^, **W**^(σ)^, **W**^(**C**)^ work as weighted adjacency matrices. Furthermore, eigenvector of last mode of *transition tensor* **<u>P</u>** would be fed into Hierarchical clustering. The number of clusters is defined as exactly same with the number of extracted components R. The stable component would produce a tight cluster. The stability index is quantified with the average intra-cluster similarities. Ideally, if the extraction of the component is stable, the inner similarity of the corresponding cluster is close to 1. The stability index of unstable components is approach to 0. The algorithm stability is defined as the average of components stability indices. When the selected model order is appropriate to the tensor to be decomposed, the algorithm would also be stable. Hence, the hyperparameter such as model order can be recommended in terms of algorithm stability.

## Notes

### Competing Interest Statement

The authors have declared no competing interest.

